# Pulsed sounds caused by internal oxygen transport during photosynthesis in the seagrass *Halophila ovalis*

**DOI:** 10.1101/2024.05.14.594143

**Authors:** Hin-Kiu Mok, Yen-Wei Chang, Michael L. Fine, Keryea Soong, Yu-Yun Chen, Richard G. Gilmore, Linus Yong-Sheng Chiu, Shi-Lin Hsu, Hai-Jin Chang

## Abstract

Oxygen bubbles that leak from seagrass blades during photosynthesis have been hypothesized to cause cavitation sounds in aquatic plants. Here we investigate low-amplitude sounds with regular pulse rates produced during photosynthesis in seagrass beds of *Halophila ovalis* (Qitou Bay, Penghu islands and Cigu Lagoon, Taiwan). Sound pulses appear in the morning when illumination exceeds 10,000 Lux, peak at midday and decrease in midafternoon on a sunny day. Frequencies peak between 1 to 4 kHz, durations range between ca. 1.8 to 4.8 ms, and sound pressure level 1 cm from the bed is 80.3±2.0 dB re 1μPa (1-3 kHz band, 11:00 on a cloudy day). Sounds attenuate rapidly, disappearing beyond 15 cm. Blocking sunlight or administering herbicide stops ongoing sounds. Gas bubbles are not typically seen during sound production ruling out cavitation, and external force (finger pressing or scissor cutting) applied to the substrate of seagrass patch or leaves, petioles, or rhizomes generally increases pulse rate. We suggest sound emission is caused by internal oxygen transport through pores in diaphragms (a whistle mechanism) at the leaf base and nodes of the rhizome.

## INTODUCTION

Seagrasses have aerenchyma tissues with continuous air-filled lacunae in leaves, rhizomes, and roots (Armstrong, 1979; Larkum et al., 1989; Borum et al., 2006; Mckenzie, 2008). The lacunae allow movement of photosynthetic oxygen produced below the leaf epidermis to flow to petioles, stem, rhizomes, and roots (Roberts et al., 1984; Lee et al., 2023) by phase diffusion (Sorrell and Dromgoole, 1987). The lacunae in leaf and rhizome are connected and have diaphragms at the nodes and transitional regions (Larkum et al.,1982). The diaphragms are perforated, containing 0.5-1.0 μm interstitial pores (Roberts et al., 1984). Seagrasses, unlike terrestrial angiosperms, lack stomata (Roberts and Caperon, 1986), thereby preserving oxygen although some is lost by diffusion through the thin cuticle.

During high levels of photosynthesis, increased oxygen pressure in the lacunae can lead to a doubling of the leaf volume, and a continuous stream of bubbles will escape from cuts on the seagrass surface (Roberts and Caperon, 1986). Likewise, turtle grass *Thalassia testudinum* leaves swell, doubling above their early morning volume, due to production of photosynthetic gases (Zieman, 1974). He commented that “by early afternoon on a shallow calm *Thalassia* bed with little water flow, the bed can readily be heard to hiss from the rapid bubbling” and “gave the appearance of a “newly open: bottle of beer.” Further, Felisberto et al. (2015) noted ambient noise from a *Posidonia oceanica* seagrass bed exhibited a diurnal pattern with most energy between 2-7 kHz.

Borum et al. (2006) determined that the largest loss of oxygen from seagrasses is from leaves to the water column during periods of high light. The continuous leakage of oxygen from roots and rhizomes to the anoxic sediment both during light and dark periods also represents a major oxygen sink. As substantial production of O_2_ occurs during photosynthesis, Borum et al. (2006) suggested that the bursting stream of bubbles emerging from the seagrass leaf may be responsible for producing sounds and that a passive acoustic system could be used to monitor the O_2_ production of a seagrass meadow. During photosynthesis in submerged waterweed, *Elodea canadensis*, gas bubbles are generated, and a series of sound pulses can be produced when the bubbles exit damaged tissue caused by the physical force of waves, currents, or propellers, etc. However, these sounds are not caused by the passage of the bubbles in the water (Kratochvil and Polliver, 2017). The pondweed *Potamogeton* spp., generates ultrasonic popping sounds with energy between 18-25kHz via leaking gas bubbles from the stomata (Kratochvil and Polliver, 2017; Greenhalgh et al., 2023). At high but unspecified temperatures, harmonic sounds with continuous frequency changes often occur (Kratochvil and Polliver, 2017). Kratochvil and Polliver suggested they are caused by internal gas flows. In seagrass, mass flow of oxygen could theoretically occur on a small scale driven by internal pressurization from photosynthesis or by leaf movement due to external physical force (Borum et al., 2006). However, no one has reported specific sounds generated by such movement.

During an acoustic survey of spawning aggregations of sciaenid fishes in the Indian River Lagoon, Florida, U.S.A. (Mok and Gilmore, 1983), we recorded continuous series of click sounds on sunny days in meadows of Manatee Grass, *Syringodium filiforme* and Shoal Grass, *Halodule wrightii*. We find similar sounds in meadows of other seagrasses on the Southwestern coast of Taiwan, Penghu Island and Dongsha Atoll in the South China Sea (Mok, unpublished data).

In this study we describe the sounds produced by *Halophila ovalis* (Fig. 1A), their diurnal periodicity, test the effect of light intensity, block light and expose seagrass to a photosynthetic blocker inhibiting sound production. We hypothesize that pulsed sounds are caused by flow of oxygen gas in the lacunae, which passes through the pores on the diaphragms and test this hypothesis experimentally.

**Fig. 1.**
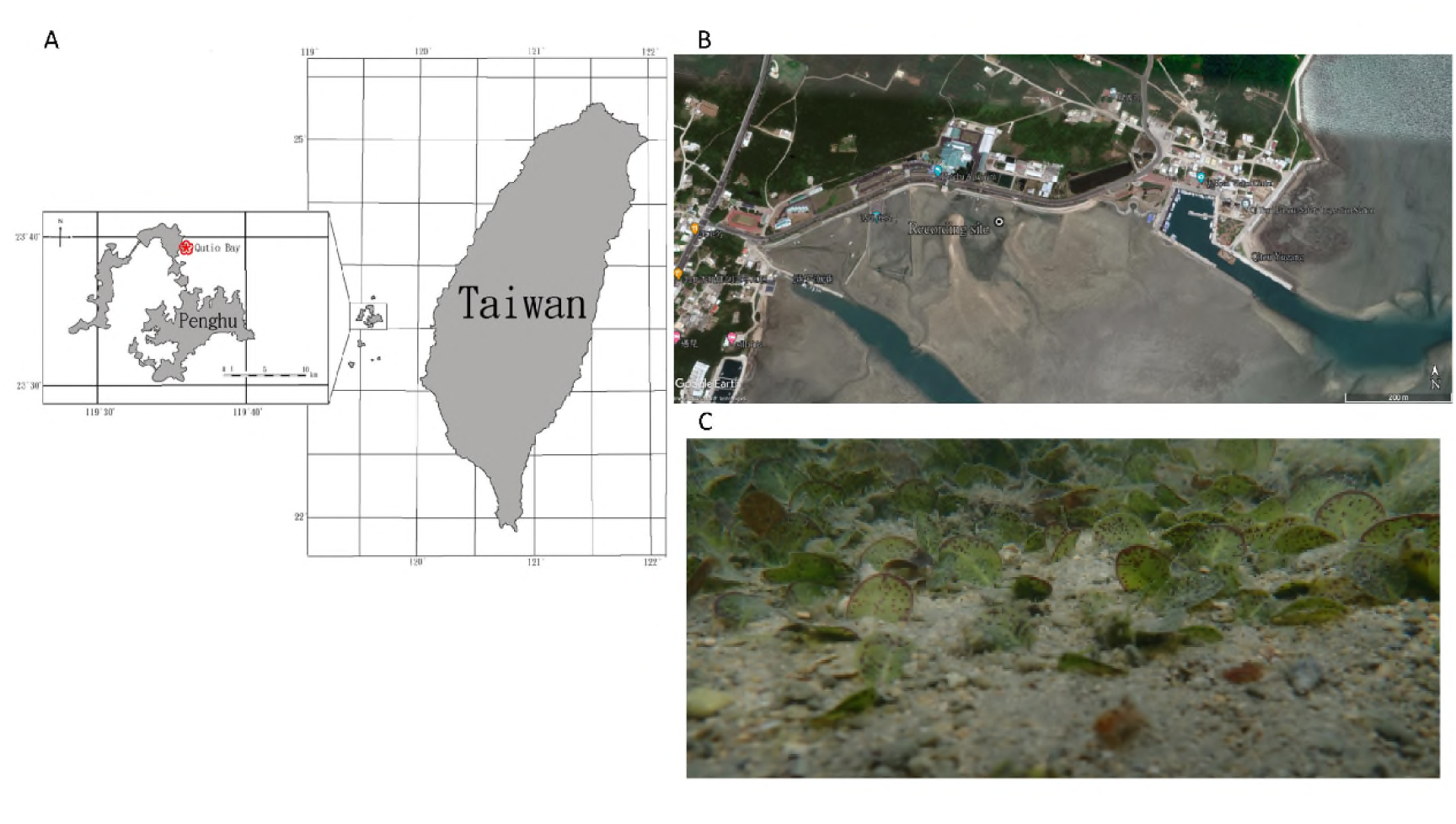
Locality of the study site Qitou Bay, Penghu islands. (A and inset), and the close-up view of the bay (B). C, In-situ view of *Halophila ovalis* patch at the bay.

## MATERIALS AND METHODS

Dugong (spoon) grass, *Halophila ovalis* (Hydrocharitaceae) was studied because of its small size compared with other seagrasses in Taiwanese waters and its accessibility for experimentation (Fig. 1A). It occurs in the Indo-Pacific region (Kuo, 2020) and is abundant in Qitou Bay (23°38′55″ N 119°36′19″ E) Penghu Islands (Fig. 1B, C).

Diel ambient sound was monitored on 24-25, July, 2020 and 25, August, 2021. Depth at the recording site at high tide was ca. 2 m and about 10 cm at low tide. Additionally, experiments were conducted at Qitou Bay and at Cigu Lagoon (23°09′18.2″N 120°06′18.4″E) on the southwest coast of Taiwan during low tide when the seagrass patches were submerged at depths < 10 cm (also see below).

Pulse series and its composing pulses are sorted based on pulse rate and power spectral density level. Hourly changes in pulse-series types, pulse rate and pulse number in daytime were estimated by sampling hourly from recordings between 0600 to 1800hrs on 07/25/2020. Pulse rate was determined from four 5-sec sections from the first 5 min in each quarter hour, e.g. 16 sections per hour. Means were and standard deviation were used to estimate pulse number per hour. Pulse duration was measured for the first six cycles in the pulse because subsequent cycles decrease in amplitude and become hard to recognize.

Sounds were recorded from six patches of *H. ovalis* and six control sites in nearby sandy areas without vegetation at Qitou Bay, and amplitude was measured at Cigu Lagoon during low tide in sunny conditions and at night. The sound pressure level was estimated from a patch of *H. ovalis* with a calibrated system placed on the substrate ca. 1 cm from the sides of the leaves.

At the *H. ovalis* meadow at Qitou Bay from 07/24-25/2020, illumination intensity ranged from 0-103,137 lux, and temperature ranged from 27.2-42.5℃. Maximum temperature occurred at 11:26hrs, and the period of high temperature (above 40℃) lasted from 09:41-11:29hrs. High tides occurred at, 13:08, 8:06 and 13:57hrs, and low tides at 19:34, 08:06, and 20:20hrs.

To test if sound emission is related to photosynthesis, three patches of seagrass were collected, and substrate sediment and invertebrates were removed. Samples with wet weights of 121g, 110 g and 132 g were placed in separate plastic trays (45 x 35 x 12 cm) filled with seawater to a depth of ca 10 cm. The trays were placed on the shore in close proximity, and a hydrophone was placed on each patch. Light intensity and temperature were recorded in one tray. Sound was monitored continuously during four 15 min treatments: control (normal sunlight condition); trays covered by a transparent plastic plate, by a black plastic plate (dark condition) and then uncovered, a return to normal sunlight. Finally, we added a photosynthesis inhibitor (Diuron dissolved in absolute alcohol to the seawater at 0.01 g per liter) and recorded sounds for 30 min.

To evaluate the relationship between sound pulse emission rate and light intensity, a patch of *H. ovalis* was placed in a 29 cm x 17 cm x 24 cm aquarium with 10 L of artificial seawater. A plant grow-light set (Smart Grow Light, SJZX-ZH001; with 8 brightness levels) faced the front of the aquarium 30 cm from the HOBO behind the tank. Sounds were recorded for five min at 5 brightness levels, 10%, 30%, 50%, 70% and 100%; the recorded illumination intensity at 100% was ca. 18,200 lux.

To test if sounds are produced by oxygen bubbles leaking from the seagrass or floating to the surface, a hydrophone was placed inside a beaker with a cleaned seagrass patch, and a video was recorded. The same procedure was conducted in the field. The beaker test was conducted in this one case because bubbles were seen escaping from the plant.

To test if sound emission and acoustic characteristics are related to oxygen transport from the leaves to the rhizomes and roots, the following experiments were conducted.

(1). Trays were monitored by video camera, hydrophone, and light intensity gauge in the morning as sunlight increased. To monitor potential bubble release during photosynthesis activity at dawn (e.g., 06:30 to 08:00), a video camera was placed adjacent to the hydrophone and the seagrass patch. Similar video and audio monitoring was also conducted on undisturbed plants in the field.
(2). Sound was monitored in an intact patch of seagrass in the meadow in which leaf was compressed by fingers and the patch was pressed by hand.
(3). A hydrophone was placed on a patch of seagrass in Qitou Bay to record sounds for 5 min after which individual leaves were cut slowly and transversely from the petiole at the base of the blade. The detached leaves were left in the vicinity of the petiole, where any sounds would be recorded. This procedure was repeated three times.

### Recording systems

We used two Sony PCM-M10 digital recorders (frequency response 20 Hz to 20 kHz), and one Sony PCM-A10 High-Resolution Audio Recorder (frequency responses linear PCM 44.1kHz, 24Bit, from 40 Hz-20 kHz). Recorders were connected to hydrophones (Aquarian Audio H2A, frequency range: <10 Hz to >100 kHz). A third recorder was placed inside a water-tight housing. Finally, a calibrated recording system including an Aquarian Audio H2A hydrophone connected to a Sony D100 PCM Recorder (linear PCM 192 kHz, 24 Bits) was used to measure sound pressure level.

For continuous recording of illuminance and temperature, a HOBO pendant MX temperature/light data logger (MX2202; temperature: 20℃ to 70℃; light: 0-320,000 lux) was used.

Diel changes in sounds from an *H. ovalis* patch at Qitou Bay, were tracked with the water-tight housing recording system and a HOBO for temperature and light data for 24 hours

Pulsed sound analysis: Avisoft-SAS Lab Pro 5.2.08 and Matlab were used for signal analysis. The recorded wave files (sampling rate at 44.1 kHz) were down-sampled at 12 kHz. Acoustical parameters including the pulse rate (number of pulses per second, in Hz, for those pulse series with a homogenous pulse rate), number of pulses per 1200s, inter-pulse interval, power spectral density, spectral centroid (which measures the spectral position and shapes and considered as the center of ‘gravity’ of the spectrum) and peak amplitude (mV) were measured from oscillograms, spectrograms and power spectra. Sound pressure levels were measured using Raven pro 1.5.2. Temporal spectral changes in underwater soundscapes were analyzed using a software assessable at (https://colab.research.google.com/drive/1Kd94iI2WjhT8KeGSQx4FeDdaD7o_gjlU? authuser=1).

### Statistical analysis

T-test (Socsci.statistics,com/tests/studentttest/default.aspx) and ANOVA (http://statskingdom.com) were conducted to test for significant differences (p<0.05).

## RESULTS

### A. Pulsed sound characteristics

Pulsed series with regular pulse rates occur primarily during the day and decrease in midafternoon and then more markedly in late afternoon and evening (Figs. 2, 3). In evening and early morning (0500 and 0600hrs), pulses occur irregularly.

**Fig. 2.**
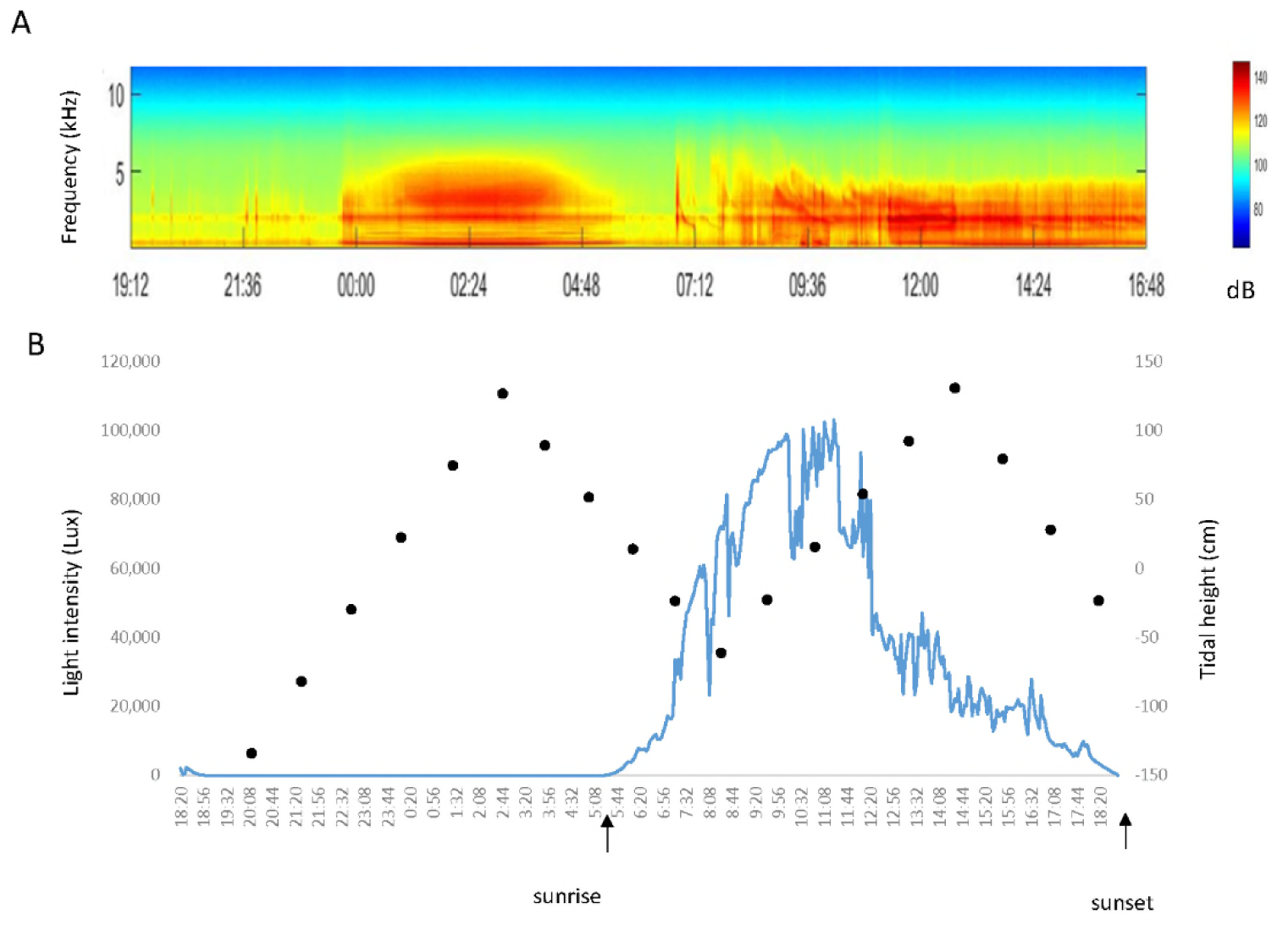
Spectrogram showing diel change in power spectral density during daytime of 20/07/24-25. (A) and temporal change in sunlight intensity and tidal height (black dot). Sunrise and sunset times at 0529hrs and 1848hrs, respectively. Loud pulses occurring from midnight to early morning (ca. 0500hrs) are irregular in pulse rate and their mechanisms remain unknown.

**Fig. 3.**
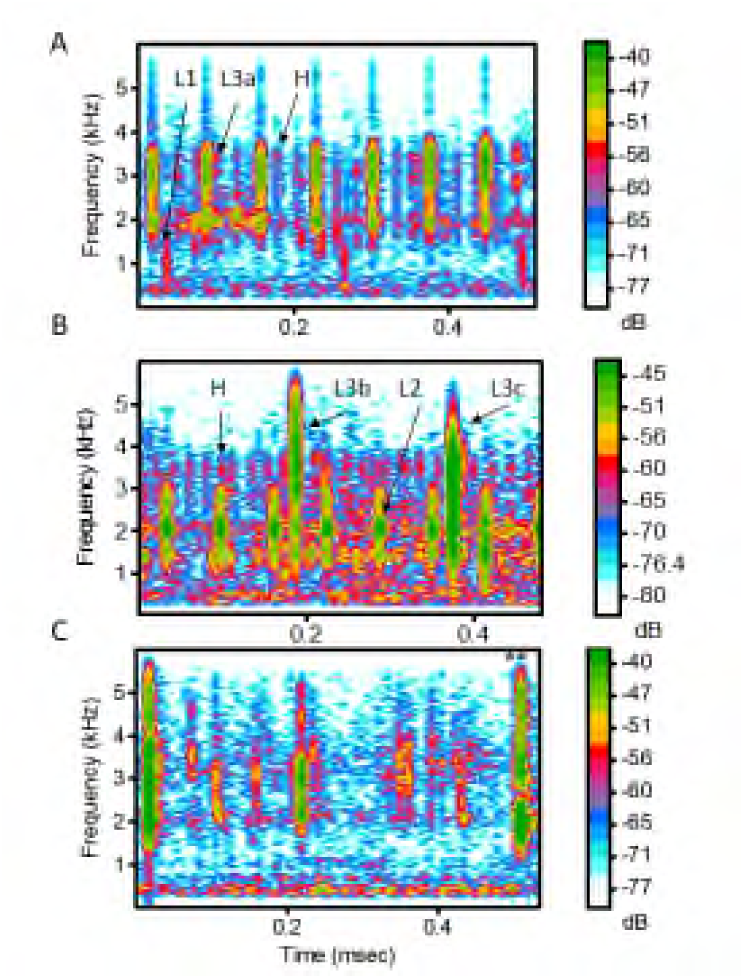
Three spectrograms of the pulsed sound from the *Halophia ovalis* patch recorded at 1000hrs. (A), 1200hrs (B) and 0200hrs (C) showing arrangement of pulse series with different pulse rates and spectral densities. Arrows point to pulse types of which the waveforms and power spectra are shown In Fig. 3. FFT=256, Frame length 100, overlap 88%. H: High-rate pulse; Low-rate pulse includes L1: Acoustic energy mainly <than 2Khz; L2: Acoustic energy mainly distribute around 2kHz; L3a: energy broadly spread between ca. 500 Hz to 4.5kHz, with energy peaks at ca. 2.0 and 3.0 kHz; L3b: with low energy bands in its frequency range; L3c; without obvious peaks in the frequency range. ** Oscillogram, power spectrum and spectrogram of the labelled pulse occurred at night are illustrated in Fig. 4G.

**Fig. 4.**
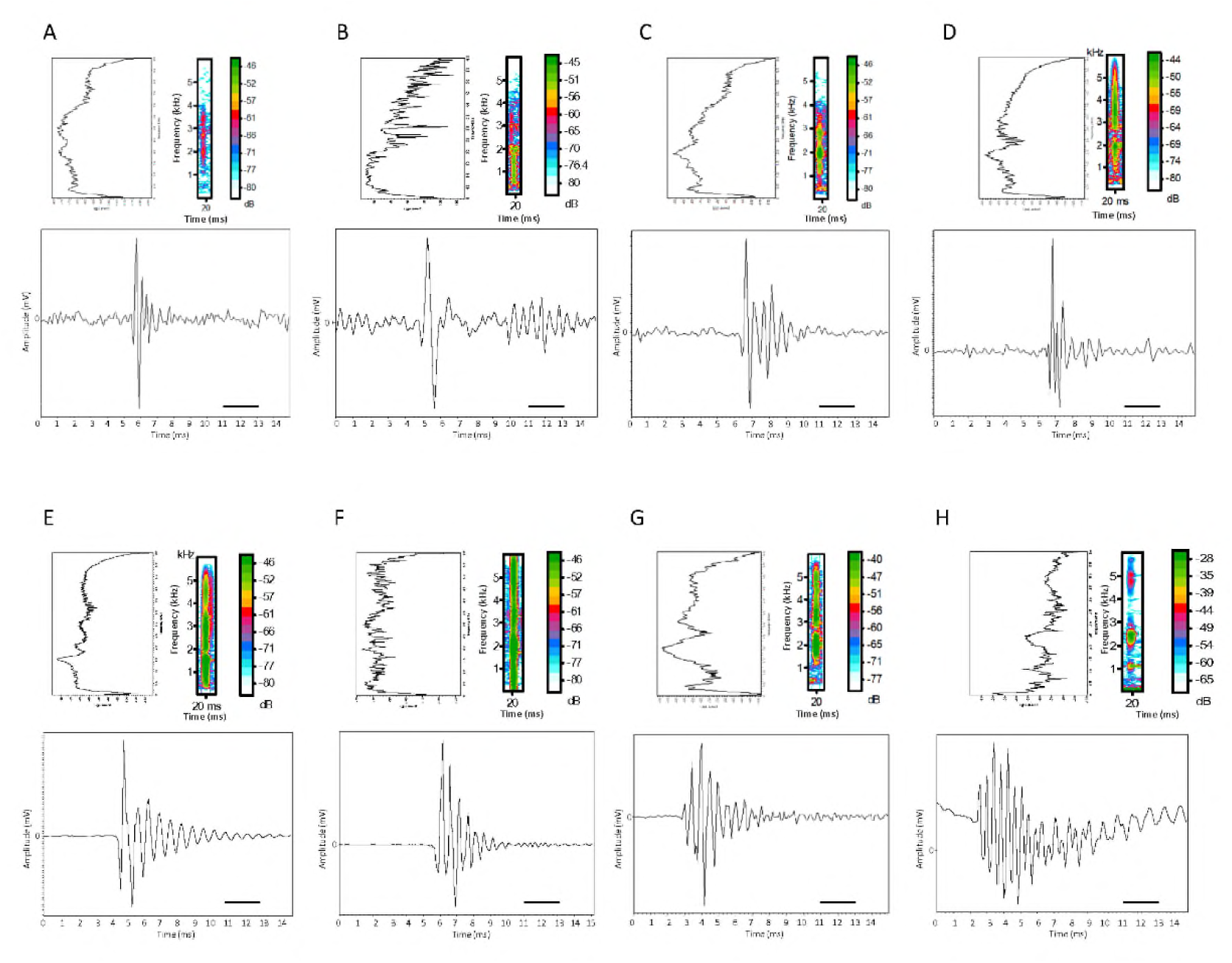
Oscillogram, spectrogram, and power spectrum of three main pulse types (high-rate and low-rate pulse types) as described in. Fig. 3. A, H; B, L1; C, L2; D, L3a; E, L3b; F: L3c; G: Pulse occurring at 0200hrs (indicated by ** in Fig. 3C); H, pulse associated with bubbles leaking out from *Halophila ovalis* in a beaker. Scale bar=2.0ms.

Series of pulses with stable pulse rates occur between 0700 to 1700hrs. They were sorted into high-rate and low rate series, pulse rates respectively ca. >38 pulses/s and <18 pulses/s; Fig. 3). Low pulse rates can be further sorted into three sub-categories by power spectral density, namely, L1, most energy below 2kHz; L2, energy peaks at ca. 2 kHz; L3, energy from ca. 0.5kHz to 5kHz. Acoustic energy in the high-rate pulses is concentrated at ca. 2kHz and 3kHz (Figs. 3, 4). Pulse durations (measured for the first 6 cycles) of these four pulse types are 3.0±0.4 ms (2.4-3.9), n=84 (H); 3.8±0.5 ms (3.2-4.8), n=38 (L1); 3.3±0.2 ms (3.1-3.6), n=30 (L2); 3.0±0.4 ms (2.4-3.9) n=84 (L3).

Temporal occurrence of series types varies (Fig. 5). The high-pulse rate series only occur between 0900 to 1200hrs, with peak at 1200hrs (Fig. 5A). In the low-rate series, sub-categories L1 and L2 appear between 0700 and 1400hrs, whereas subcategories L3 appears from 0700 to 1600hrs (Fig. 5B). Series L1, L2, and L3 peak at 1000, 1100, and 0700hrs, respectively (Fig. 5B). 1100, 1200, and 1300hrs consisted of more low-rate pulses than other hours of the daytime period (Fig. 5C).

**Fig. 5.**
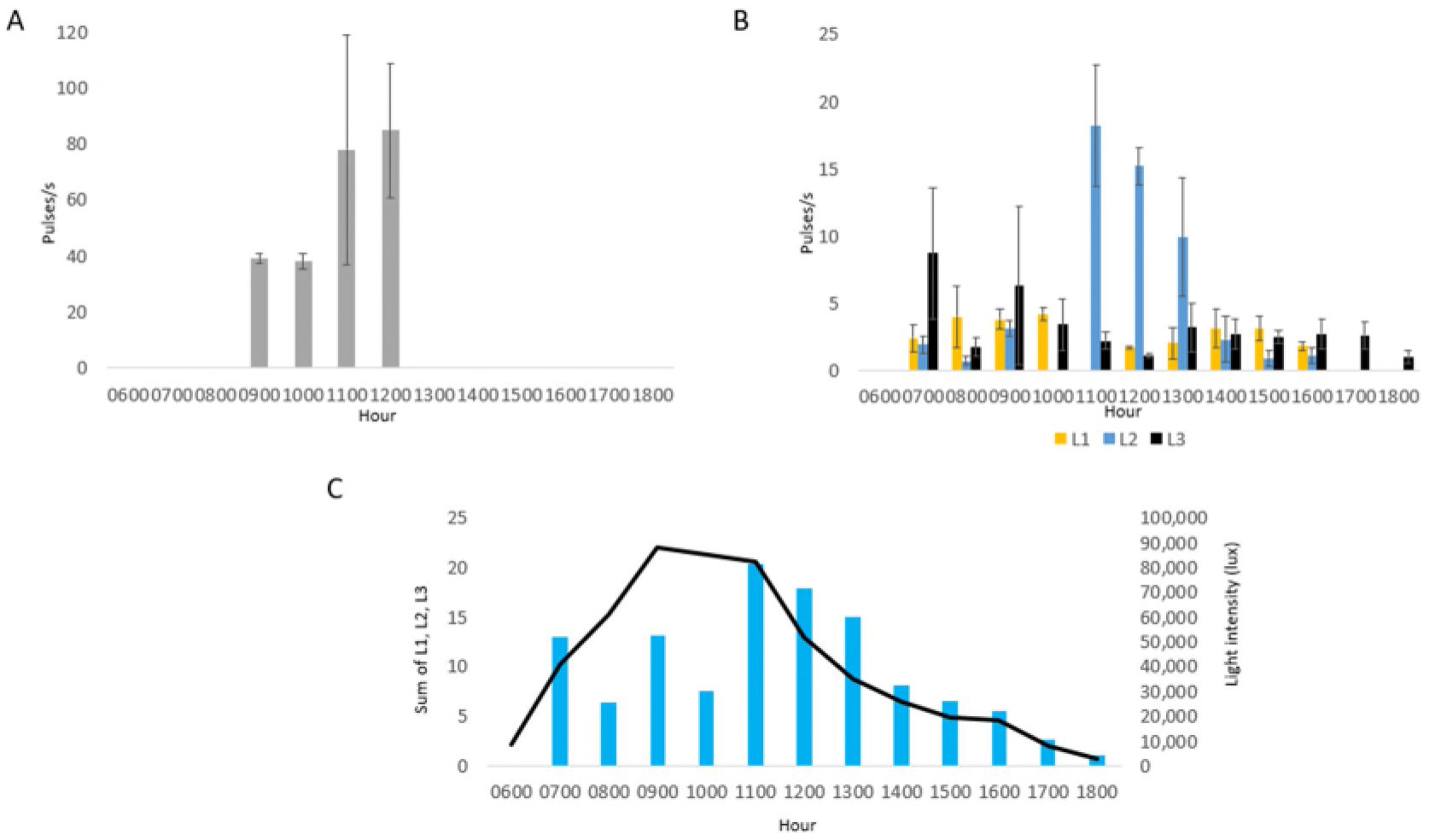
Hourly occurrence of the pulse types in the daytime shown in bar charts. A, Type H; B, L1, 2, 3; C, Average pulse rates and average sunlight intensity (line). H: high-pulse rate series; L1: low-pulse rate series, with most energy below 2kHz; L2: low-pulse rate series, with energy spread adjacent to 2kHz; L3: low-pulse rate series, energy spread from 0.5 to ca. 5kHz, with variable power spectral density, or peaks at ca. 2 and 3kHz.

On sunny days, a patch in Qitou Bay produced series of high-amplitude broad-band pulsed sounds (500-6000Hz) with stable inter-pulse intervals (e.g., 6 pps) mixed with weaker but high-repetition-rate pulses (as much as 89 pps) with peak frequencies at 2000 and 3000-4200Hz (Fig. 3). Duration of high-amplitude pulses was 3.4±0.3ms (range: 2.9-4.5ms; N=80) compared to 2.8±0.7ms (range: 2.0-3.2ms; N=80) (t_158_=6.1574; p=0.00001) for the weaker high rate series. In some instances, only series with pulses at different repetition rates occurred. During periods with bright light, pulse rates ranged from 16 to 89 pps; and a high pulse rate series could last for 200 seconds before tapering and then stopping. Daytime pulses, increase from 1 to ca. 90 pulses per sec with light intensity and often have a regular pulse rate. Maximal pulse rate occurred between 1200 and 1300hrs (Figs. 5, 6; Table 1).

**Table 1.** Spectral centroid and difference between the amplitudes for the dominant-frequency peak at ca. 1960Hz and secondary frequency peak at ca. 2900Hz.

| Time of day* | Spectral centroid (Hz) | Amplitude (dB) at ca. 1960Hz higher<br>than that at 2700Hz |
| --- | --- | --- |
| 18:20-19:20 | 2830 | 7.1 |
| 19:20-20:20 | 2850 | 6 |
| 20:20-21:20 | 2900 | 6.8 |
| 21:20-22:20 | 2620 | 7.7 |
| 22:20-23:20 | 2560 | 8.1 |
| 23:20-00:20 | 2530 | 7.4 |
| 00:20-01:20 | 2670 | 6 |
| 01:20-02:20 | 2820 | 2.5 |
| 02:20-03:20 | 2800 | 2.9 |
| 03:20-04:20 | 2810 | 2.1 |
| 04:20-05:20 | 2740 | 2.6 |
| 05:20-06:20 | 2630 | 6.1 |
| 06:20-07:20 | 2870 | 5.4 |
| 07:20-08:40 | 2660 | 8.4 |
| 08:40-09:40 | 2600 | 6.9 |
| 09:40-10:40 | 2510 | 2.1 |
| 10:40-11:40 | 2430 | 5.6 |
| 11:40-12:40 | 2440 | 5.1 |
| 12:40-13:40 | 2210 | 8.1 |
| 13:40-14:40 | 2270 | 8.3 |
| 14:40-15:40 | 2270 | 8.2 |
| 15:40-16:40 | 2240 | 6.4 |
| 16:40-17:40 | 2470 | 4.7 |
| 17:40-18:40 | 2530 | 3.9 |

\* Measured from the long-term (one-hour) average spectra from 18:20/24/07/2020 to 18:20/25/07/2020. Frequencies become lower in high light levels.

Pulse series with a single peak at 1.4 kHz diminished at ca. 15:30hrs when the light intensity was 18,304 lux. Between 16:30 and 18:20hrs energy decreased at the dominant frequency peak (ca. 1930 Hz) —the main energy peak for photosynthesis pulses (see below; Fig. 5; Table 1).

Pulses with two different rates sometimes appear simultaneously during the day at high light levels (Fig. 3). Series can start with a high pulse rate and then decrease or vice versa without changes in light intensity (see below), suggesting variable rates of photosynthetic gas production or a lag between light intensity and oxygen production. Irregularly-paced pulses also occur at night (Figs. 3C, 4G), e.g., 01:00 to 02:00; ca. 611.9±123.2 pulses per 5 min; range ca. 443-804 pulses; N=7). Those occurring between 01:00 and 04:00 contributed high acoustic energy compared to other parts of the evening and early morning (Fig.6; Table 1) are had a higher spectral centroid located between ca. 2700 and 2800Hz (Table 1), and a bigger energy difference occurred (i.e., 2.1 to 2.9 dB) between the dominant and secondary frequency peaks (ca. 1930 and ca. 2700Hz, respectively). The pulses at this period are relative high pitched (Fig. 6; Table 1).

**Fig. 6.**
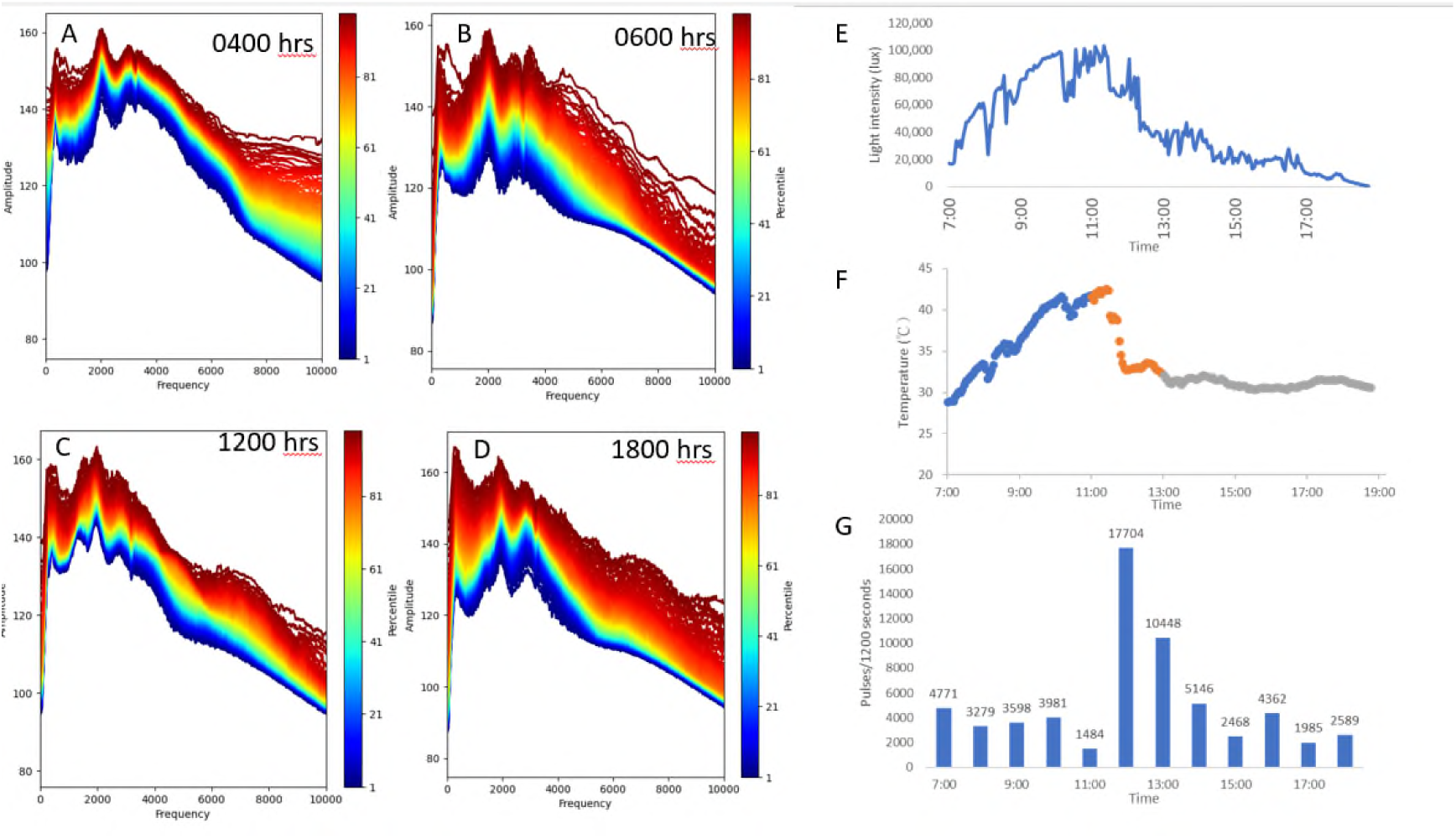
Temporal variation in power spectral range in 4 hourly sections A to D and variations in light intensity, temperature, and pulses/20 min (E to G). Notice the higher amplitude and broader range at the frequencies above 7 kHz at 0400 and 1800hrs A, D).

Waveforms are variable; the first six pulses in all the pulse were relative easy to recognize. The second cycle is often but not always weaker than the following cycles (Fig. 4). The amplitude rises abruptly with a peak at the first, second, or third cycle, followed by variable decay (Fig. 4). However, in most of the pulses analyzed from a sunny day, the first cycle has the highest amplitude and the second cycle is often weaker than the adjacent cycles (Fig. 4). Pulses with energy more homogeneously spreading across the frequency range show a smooth decay in amplitudes (Fig. 4A, F). In night time, early morning and late afternoon pulses, the second or third cycle reaches the maximum amplitude of the pulse (Fig. 4G), whereas the first cycle in the pulses under bright light has the maximum amplitude (Fig. 4A-F).

The mean sound pressure levels (SPL) of a high-amplitude pulse series with a pulse rate of ca. 14 pps at Cigu Lagoon was 100.2 ±1.9 dB re 1 µPa (range: 75.6 – 84.8 dB; N=30) in the frequency band between 1 and 2 kHz and 17 dB above background (80.3±2.0 dB, N=6), (hydrophone 1 cm from the seagrass patch, light intensity 50,000 to 60,000 lux on a sunny day and low tide). SPL for the signature frequency peak at 2.1 kHz was 83 .0±2.7dB. The sound attenuated rapidly over open sandy areas, and amplitude decreased by 17 dB at 15 cm; No signal was recorded at 20 cm from 3 patches (Fig. 7). The pulsed sounds can be barely heard when one’s ear is next to the seagrass patch (pers. obs.).

**Fig. 7.**
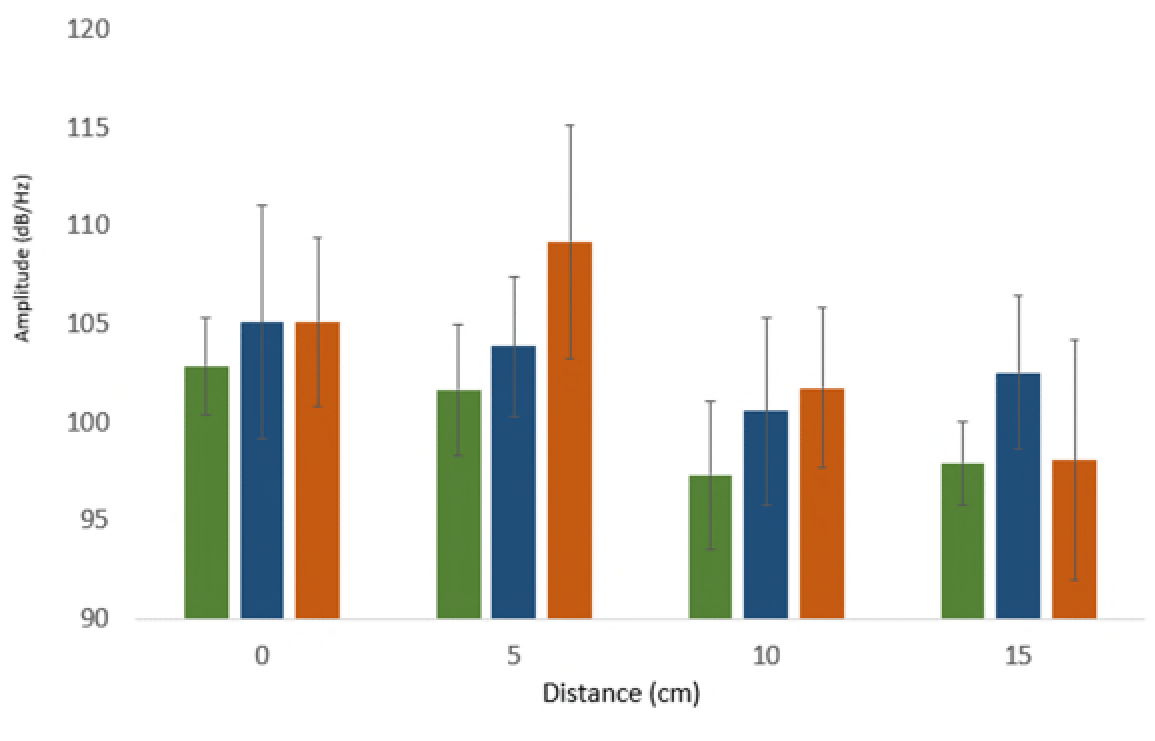
Attenuation of amplitude (mean and SD) of pulsed sounds from the edge of *Halophila ovalis* patches. Three replicates, each with 30 measurements.

### B. Experiments on pulsed sound production due to photosynthesis

#### Sound and light intensity

In samples in plastic trays under bright sun, pulsed sounds with stable inter-pulse interval were present, indicating functional photosynthesis on carefully removed plants. These sounds rule out the possibility of production by marine invertebrates cryptically associated with the rhizomes and roots.

Pulse rates in bright direct-light (50,000 and 68,000 lux) did not change when trays were covered by a transparent plate (401±72.7 and 402±46 pulses/5 min, N=3). Pulses immediately vanished after the tray was covered with an opaque black plastic plate and reappeared ca. 90 seconds after plate removal. Pulses then regained their normal repetition rate in ca 173 s. Therefore, light is required for pulse production, and pulse rate increased with light intensity (Fig. 8).

**Fig. 8.**
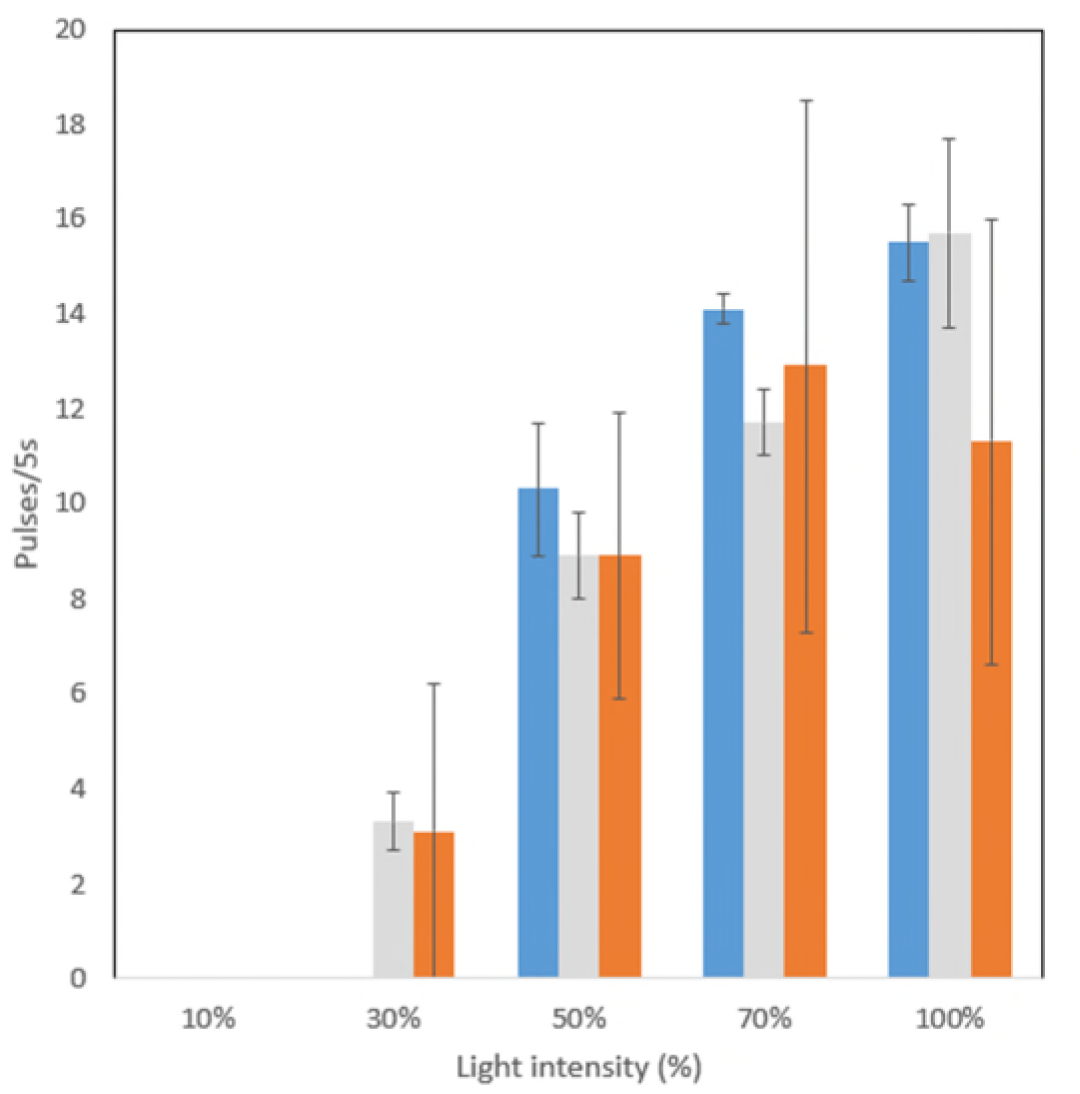
A bar chart showing relation between number of pulses/5s at 5 brightness levels in three trials tested in an aquarium. Illuminance at 10%, 30%, 50%, 70%, and 100% were 790, 3970,7440,10270, and, 16870 lux, respectively. Notice absence of pulsed sound at the 10% brightness level.

In emission tests on an intact seagrass patch at Cigu Lagoon (60,000 and 70,000 lux in light and 80 Lux in dark), pulse rate decreased from 298.7±43.7 pulses/30s (N=6) to 0 pulses in less than 1 min when covered. A replicate on a near-by patch likewise decreased similarly from 277.5±39.8 pulses/30 s (N=6) to 0 pulses/30 s.

Pulse rate decreased 30 s after Diuron addition and vanished completely after about 23 minutes in three trials. Light intensity at the bottom of the plastic container varied between ca. 50,000 to 60,000 lux.

#### Pulsed sounds and bubbles leaking from the seagrass

On one occasion under sunlight conditions, we saw gas bubbles emerging from a small patch of *H. ovalis* placed in a beaker. Pulse rate was 11.8±3.4 pulses/5s or 2.4 pulses/s (N=14). Pulse duration was 11.1±1.4 ms (range 10.1-15.9ms), with three pronounced frequencies at ca. 1.0, 2.5, 5 kHz (Fig. 4H). Maximum amplitude of a pulse occurred in the third cycle (Fig.4H). Therefore, gas bubbles leaking from the seagrass can generate sounds, but they were different in duration, spectral density and waveform from typical photosynthesis pulses (i.e., longer duration, more cycles per unit time, and amplitude peaked at the third cycles instead of the first cycle; Fig. 4H). Streams of minute air bubbles were noted during brightness testing in an aquarium, but no sounds were recorded adjacent to the bubble streams.

#### Sound emission and acoustic characteristics are related to oxygen transport in the air-channel system of H. ovalis

In cloudy conditions, when pulses had an irregular emission rate, force applied by fingers to a patch and its substrate immediately increased pulse repetition rate and peak frequency followed by a decrease. Sounds were composed of high and low amplitude pulses with variable repetition rates. High amplitude pulses had repetition rates of 44.9±22.2 pulses/s (range 18 - 87 pulses/s; N=15) vs 155.7±45.6 pulses/s (range 89 - 230 pulses/s; N=7) for low amplitude ones. Sound emission sometimes changed from a mixed series to one with only low amplitude pulses suggesting that high-amplitude pulses require greater leaf pressure. Duration of pulse series averaged 19.9±20.2ms (range 6.8 to 32.9 s; N=11).

Cutting a petiole also increased repetition rate (Fig. 9) likely because the force applied generated gas movement inside the lacunae. Pulses from a seagrass patch decreased as the leaves were cut below the leaf-petiole junction or from the middle of the leaf one by one. No pulses were present after all leaves were cut even though they remained in the vicinity of the seagrass patch. Repeated tests conducted in the trays and in the field were similar.

**Fig. 9.**
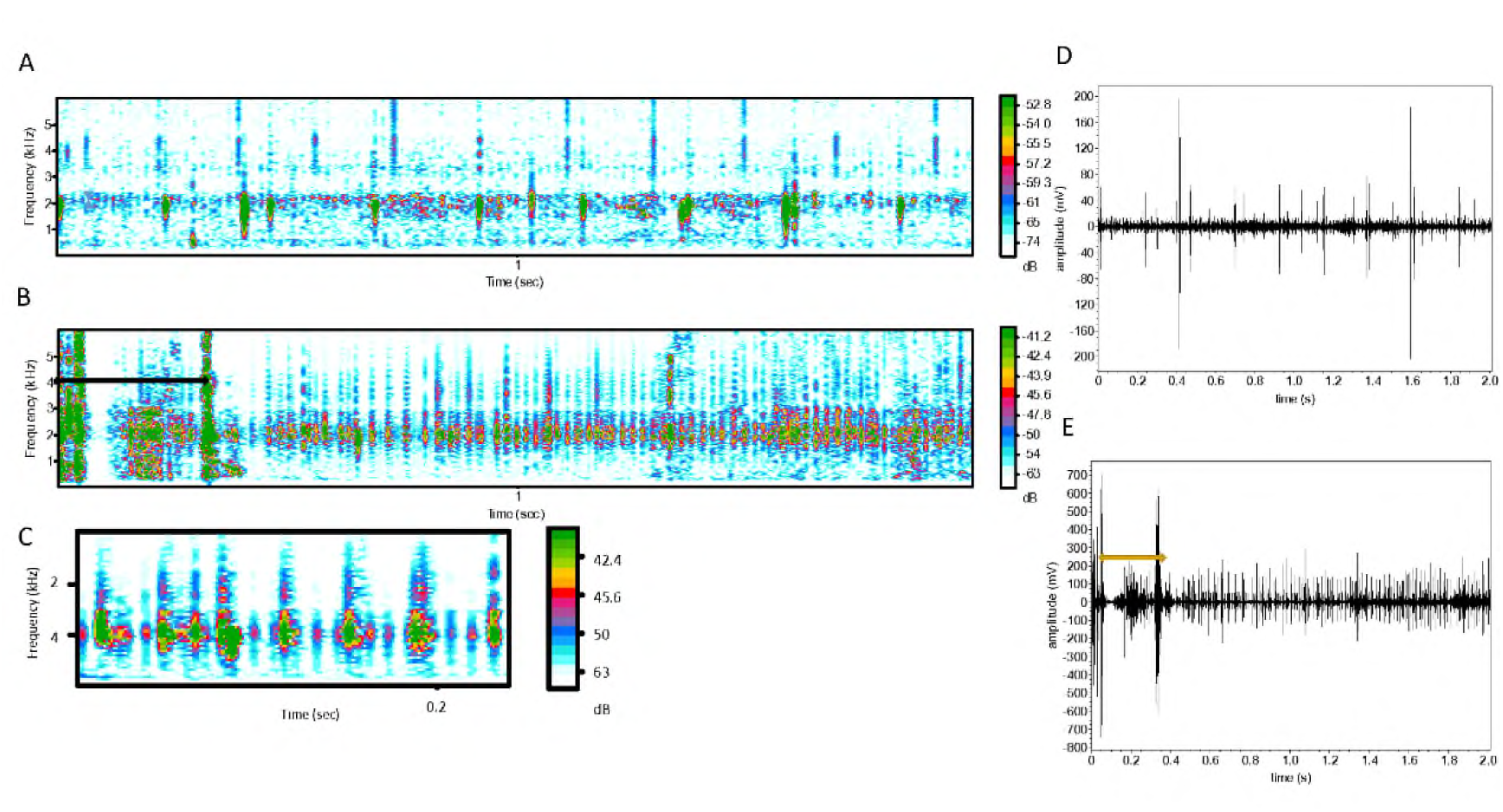
Spectrograms (A, B, C) and oscillograms (D, E) showing the changes in pulse types and pulse series prior to (A, D) and during cutting (B, C, E) the *Halophila ovalis* petiole. FFT 256 in A, B and FFT 128 in C; frame size 100 and overlap 88%; Hamming window. Double-ended arrows in B and E represent the time when the plants were cut with scissors.

## DISCUSSION

Kratochvil and Polliver (2017) reported gas bubble sounds from submerged American waterweed, *Elodea canadensis* in a small tank, and sounds ceased after lights were turned off. Frequencies ranged from 20 Hz to 20 kHz, with most energy in the high-frequency range. At high but unspecified temperatures, continuous frequency changes often occurred, and the authors suggest they are possibly caused by gas flows inside the plant.

Gas bubbles with sounds emitted from *H. ovalis* in a beaker were seen once and were associated with pulsed sounds; but pulsed sounds were not recorded from stream of small bubbles from *H. ovalis* in a small tank. However, pulses were recorded repeatedly from undisturbed *H. ovalis* patches in shallow water, during which no bubbles were seen. Therefore, bubbles are not the source of these sounds, supporting Kratochvil and Polliver’s suggestion of sounds caused by internal gas transfer.

During light periods, O_2_ pressure increases within *H. ovalis* leaves and causes oxygen flow to lower parts of the plant that inhabit an oxygen poor substrate. Pulsed sounds appear in series with irregular paces when morning illumination increases above 10,000 Lux. As light intensity increases and leaves swell from gas accumulation, pulse rate increases and becomes regular. Under bright sunlight, low amplitude pulse series emerge and mix with the high-amplitude series. Under this situation, an external force applied to leaves boosts only the rate of the low-amplitude series but not the loud series, suggesting that separate threshold gas pressures are necessary to produce low and high-amplitude pulses. Photosynthetic pulses begin to decrease around 15:00 although light intensity is still high. Fewer pulses likely reflect indicate decreased photosynthesis, which might be due to photoinhibition, temperature stress, or high carbohydrate end-product inhibition of photosynthesis (Larkum et al., 2006; Azcón-Bieto, 1983; Goldschmidt and Huber, 1992). Therefore, we suggest that pulsed sounds from plant beds could be used to reveal the health of seagrass meadows and environmental stresses. Plants covered and held in dark conditions stop producing sounds, but similarly produce high-pulse rate and low-amplitude sound series several seconds after return to high luminance. These continue and are followed by the addition of the slow rate high amplitude pulses. Pressing the patch at night or in a tray covered with black plastic does not generate pulsed sound.

Cutting the seagrass petioles or pressing the patch between hands in sunlight rapidly increases pressure in the lacunae, causing gas to escape through the 0.5 and 1 µm holes in the diaphragms between the leaf and stem, thus producing a whistle-type sound (Fletcher, 1992). Fletcher notes that the basis of aerodynamic sound arises from a steady flow that becomes unstable if it is forced to change direction or to undergo shear deformation near a solid surface. In this case gas pressure from photosynthetic oxygen in relatively wide lacunae will change direction as it passes through 0.5 to 1 µm pores in the diaphragms (Roberts et al., 1984). The system is further complicated by multiple pores per diaphragm and multiple lacunae per leaf. Therefore, pressurized gas is typically required for sound emission (but see below), and we suggest higher gas pressure thresholds for loud than quiet pulses. Warmer temperatures should also cause a minor increase leaf pressure and hence sound repetition rate and amplitude since it will increase gas volume within the leaves (Charles’ Law) and therefore flow through the diaphragms.

Under high illumination, a series of pulses has a stable inter-click interval, implying pulses come from individual sources in variable acoustic states rather than multiple leaves or plants simultaneously. Termination of a series of intense pulses from a leaf would stop when pressure decreases below threshold. Subsequent intense sounds would then likely come from another leaf.

Felisberto et al. (2015) recorded ambient noise at night with most energy between 2 to 7 kHz from a *Posidonia oceanica* seagrass bed at 2 to 20 m depths. The mean noise power was negatively correlated with dissolved O_2_, and they suggest that power is correlated with photosynthetic activity, which is unlikely since photosynthesis would not peak at night. Our night recordings (0100 to 0300hrs) also indicate sounds in the typical frequency band at ca. 2-3 kHz, which is presently unexplained.

Seagrass meadows attenuate sound rapidly compared to open shallow habitats (Chang et al., 2019; Wilson et al., 2012). The gas within adjacent seagrass leaves dissipates the acoustic-energy, and spreading of the sounds from a small bundle of seagrass should be minimal because of absorption by adjacent seagrass bundles.

The evolution of plants sounds is intriguing and unexplored. The ability to produce sounds in *Halophila ovalis* shares characteristics with evolutionary exaptations, a term coined by Stephen J Gould and Elizabeth Vrba (1982) in which a functional character is later co-opted for a new use that enhances fitness. In acoustic communication the term has been invoked for the evolution of sound-producing structures in fishes (Parmentier et al., 2017). For instance a tendon that causes rapid jaw slams in damselfishes (a feeding adaptation) has been turned into a sonic structure that functions in behavior (Parmentier et al., 2007), swimbladders that contribute to buoyancy and gas regulation have become sonic organs by attaching muscles that cause them to vibrate (Fine and Parmentier, 2022), and a defensive catfish spine that can be locked to deter predators (Sismour et al., 2013) has been modified into a stridulatory organ in which ridges on the spine base are rubbed against a rough surface on the pectoral girdle, a stick-slip mechanism as in a bow on a violin (Mohajer et al., 2015). Similarly, oxygen produced in the leaves of *Halophila ovalis* causes them to swell, and the increased pressure drives gas transport through small holes in their diaphragms, pushing oxygen to underground portions of the plant in an anoxic environment. It is unknown whether these sounds provide signals to adjacent plants, which would be necessary to establish an exaptation. For now, therefore, we consider the sounds to represent an epiphenomenon related to gas transport rather than a functional acoustic signal.

## Conclusion

We demonstrate that photosynthesis increases leaf pressure driving oxygen through pores in diaphragms that separate portions of the plant. This gas movement produces a series of pulsed sounds that can vary in pulse rate, amplitude and waveform. Pulse rates are variable early in the morning but become regular and more rapid as light intensity increases. Although higher amplitude pulses and greater pulse rate appear to be caused by higher leaf pressures, much of the variation is still unexplained.

## Acknowledgments

We are grateful to Prof. T. M. Lee, National Sun Yat-sen University who gave us advice in using herbicide Diuron and logistic support for field recording. Our sincere thanks go to Prof. Adair, B. for calling our attention to the presence of the paracrytic-type of stromata in *Thalassia testudinum*, its possible role in generating sound, and the value of acoustic monitoring for pollution impacts on the seagrass meadow. Dr. C.W. Chang, National Academy of Marine Research provided the calibrated recording system to measure sound amplitude. Mrs. P. H. Chiu, National Sun Yat-sen University, assisted us in measuring the sound pressure level. Prof. K. S. Chiu, National Kaohsiung University of Science and Technology, helped with field collection. Dr. T. H. Lin, Academia Sinica, kindly provided the software he developed that uses an algorithm for source separation models to visualize temporal spectral changes in underwater soundscapes (https://colab.research.google.com/drive/1Kd94iI2WjhT8KeGSQx4FeDdaD7o_gjlU?authuser=1).

## Authors’ contributions

M-HK: discovered the sound producing phenomenon, conception, proposing, experimental design and performance, analysis and interpretation of data, manuscript writing, final approval of the manuscript; C-YW: field recording, performing experiments; F-ML: conception, manuscript writing; S-KY: conception, experimental design, data analysis; C-YY: field recording; G-RG: manuscript critical revision, C-LYS: manuscript revision, H-SL: field recording, data analysis; C-HJ: field recording and performing field experiments. All authors read, edited, and approved the final version of this manuscript.

